# Systematic analysis of transcriptional and post-transcriptional regulation of metabolism in yeast

**DOI:** 10.1101/057398

**Authors:** Emanuel Gonçalves, Zrinka Raguz, Mattia Zampieri, Omar Wagih, David Ochoa, Uwe Sauer, Pedro Beltrao, Julio Saez-Rodriguez

## Abstract

Cells react to extracellular perturbations with complex and intertwined responses. Systematic identification of the regulatory mechanisms that control these responses is still a challenge and requires tailored analyses integrating different types of molecular data. Here we acquired time-resolved metabolomics measurements in yeast under salt and pheromone stimulation and developed a machine learning approach to explore regulatory associations between metabolism and signal transduction. Existing phosphoproteomics measurements under the same conditions and kinase-substrate regulatory interactions were used to estimate the enzymatic activity of signalling kinases. Our approach identified informative associations between kinases and metabolic enzymes capable of predicting metabolic changes. We extended our analysis to two studies containing transcriptomics, phosphoproteomics and metabolomics measurements across a comprehensive panel of kinases/phosphatases knockouts and time-resolved perturbations to the nitrogen metabolism, conveying a total of 143 unique conditions. Our approach accurately estimated the change in activity of transcription factors, kinases and phosphatases and these were capable of building predictive models to infer the metabolic adaptations of previously unseen conditions across different dynamic experiments. Time-resolved experiments were significantly more informative than genetic perturbations to infer metabolic adaptation. This difference may be due to the indirect nature of the associations and of general cellular states that can hinder the identification of causal relationships. This work provides a novel genome-scale integrative analysis to propose putative transcriptional and post-translational regulatory mechanisms of metabolic processes.

## Introduction

Cells sense and react to extracellular *stimuli* with coordinated intracellular responses conveying transcriptional, protein and metabolic changes (Chubukov et al. 2014; Herrgard et al. 2008; Daran-Lapujade et al. 2007). The continuous technological advances over the last few decades has contributed to the advent of the omics era with quantitative measurements of hundreds to thousands of transcripts, proteins and metabolites across a variety of steady-state and time resolved conditions (Bodenmiller et al. 2010; Kemmeren et al. 2014; Schulz et al. 2014). The increasing accumulation of molecular measurements have provided unprecedented knowledge of the cellular molecular adaptation, nonetheless the robust identification of the regulatory interactions underpinning these changes is still a challenge (Oliveira et al. 2012; Zelezniak et al. 2014). Currently, the bottleneck has shifted from data acquisition to the development of statistically robust and computationally efficient mathematical approaches capable of providing an integrated analysis of the different types of biological data available.

Regulatory responses mediate the adaptation of many biological aspects of a cell, for example, metabolism may be regulated transcriptionally and post-transcriptionally. At present, most of the integrative analysis of metabolomics data-sets have focused on the role of transcriptional regulation (Patil & Nielsen 2005; Zelezniak et al. 2014; Gerosa et al. 2015; Oliveira, Dimopoulos, et al. 2015). Previous studies have focused on the regulatory implication of transcription-factors (TFs) to model the metabolic transition between different steady-state conditions (Gerosa et al. 2015). Moreover, these regulatory interactions may occur in the inverse direction where metabolites directly impact the activity of global cellular regulators, such as TOR1 (Oliveira, Dimopoulos, et al. 2015; Oliveira, Ludwig, et al. 2015). Nevertheless, transcript levels have been shown to poorly predict metabolic fluxes in the central carbon metabolism and that glycolytic enzymes are predominantly regulated at the post-transcriptional level (Daran-Lapujade et al. 2007; Daran-Lapujade et al. 2004; Machado & Herrgard 2014). Signal transduction by reversible protein phosphorylation is a key cellular regulatory mechanism and has been shown to modulate the glycolytic flux by regulating metabolic enzymes (Oliveira et al. 2012). Recent studies have explored the implication of phosphosites in the enzymatic activity of kinases/phosphatases (K/Ps) by integrating with metabolomics measurements and *in silico* estimated metabolic fluxes (Oliveira et al. 2012; Yugi et al. 2014). Nonetheless, data acquisition and subsequent integrated analysis of phosphoproteomics data-sets are much sparser than transcriptomics and are still lagging behind (Oliveira et al. 2012; Yugi et al. 2014).

Transcriptional and translational regulatory interactions of metabolism can, in principle, be comprehensively explored using available high-throughput data-sets and methods (Kemmeren et al. 2014; Bodenmiller et al. 2010; Oliveira, Dimopoulos, et al. 2015). However, current methods have yet to integrate gene-expression and phosphoproteomics measurement to infer regulatory interactions of metabolism. In this study, we set out to address this issue. We propose a computational approach to systematically identify putative post-transcriptional and post-translational regulatory mechanisms of metabolism (Fig 1A). To this end, we characterised the metabolomics adaptation of yeast under salt and pheromone conditions and further expanded it to consider a compendium of experimental data-sets (Bodenmiller et al. 2010; Kemmeren et al. 2014; Schulz et al. 2014; Oliveira, Ludwig, et al. 2015; Oliveira, Dimopoulos, et al. 2015), comprising a total of 143 unique conditions. Firstly, we estimated the *in vivo* activity of TFs and K/Ps. For that purpose, we considered prior-knowledge on regulatory interactions and mathematical approaches that have been developed to infer the activity status of transcription factors (Cheng et al. 2012; Schacht et al. 2014) and kinases (Casado et al. 2013; Mischnik et al. 2016) (Fig 1A). The activity of regulatory proteins is difficult to measure directly, yet provides functional information about the protein regulators involved in a cellular response. Subsequently, regulator activities were integrated with the metabolomics measurements using a machine learning approach to infer putative regulatory interactions. Our approach accurately estimates the activity status of known regulatory proteins, and identifies protein-metabolite associations capable of robustly estimating metabolic phenotypes of previously unseen conditions.

**Figure 1.**
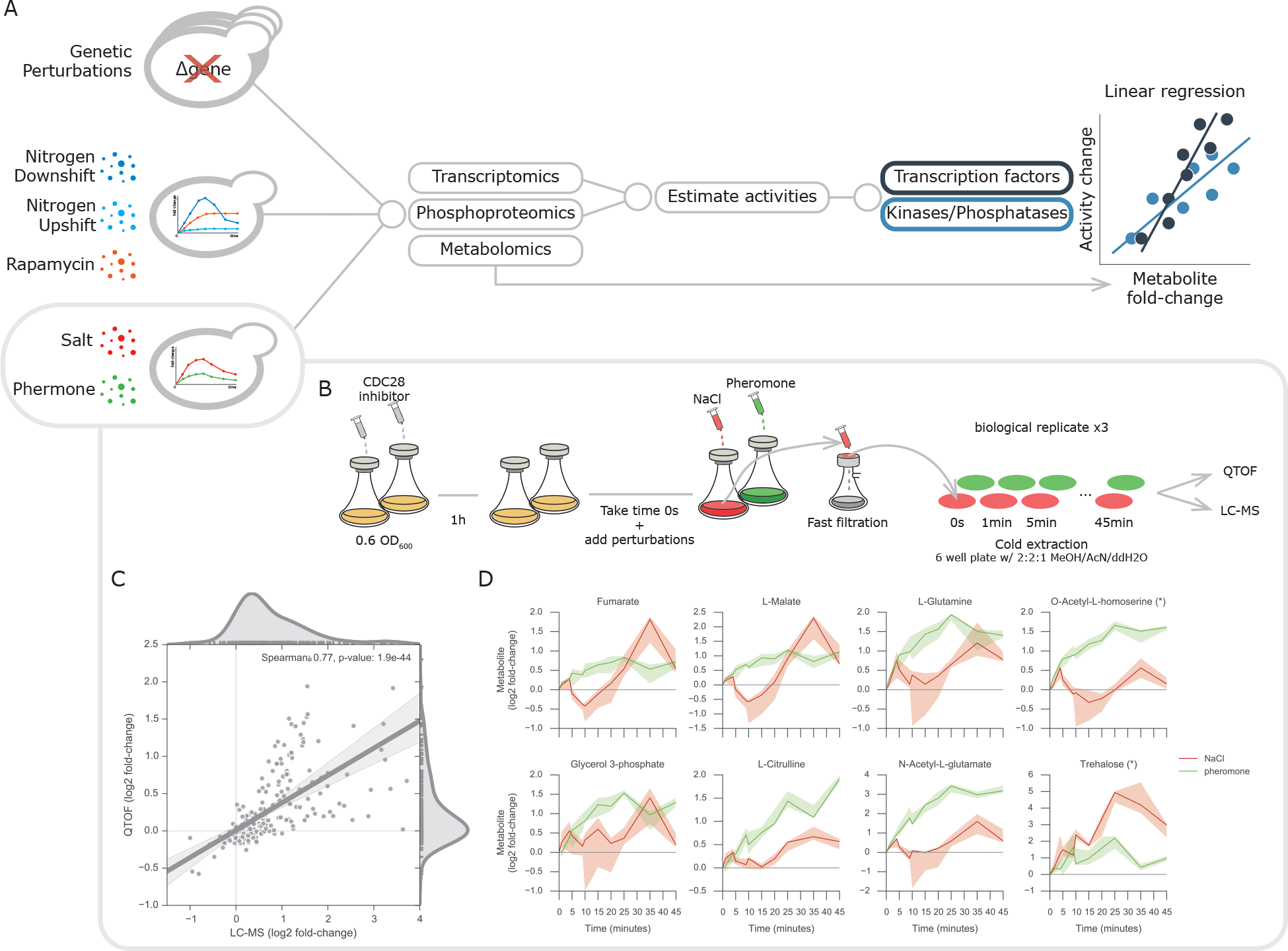
Analysis and experimental design and data consistency. (A) Representation of the different types data-sets used in the analysis. Transcriptomics and phosphoproteomics datasets are used to estimate transcription factor and kinase/phosphatase activity changes, which are then separately associated with the respective metabolomics data-set using multilinear regression models. (B) Experimental design used to acquire the intracellular metabolomics measurements. CDC28 analog sensitive yeast strains inoculated in shake flasks were treated with the CDC28 inhibitor. The unperturbed initial time points were taken 1 hour after the CDC28 inhibitor and before adding the NaCl and pheromone. Sample filtration, metabolite extraction and MS injection were performed in parallel on the samples from independent triplicate experiments. (C) Representative metabolite profiles of untargeted metabolomics experiments.(D) Metabolites fold-changes correlation between targeted and untargeted metabolomics. Ions mapping to more than one metabolite are marked with an asterisk (*) and were not considered for any downstream analysis.

## Results

### Generation of dynamic metabolomics upon salt and pheromone perturbation and integration with a compendium of existing data-sets

To explore the functional implication of post-translational regulatory mechanisms in metabolism we set out to obtain paired phosphorylation, expression and metabolomics data under the same experimental conditions. For osmotic and pheromone conditions we resorted to existing dynamic phosphoproteomics measurements and we experimentally determined the intracellular metabolite changes. Both salt and pheromones are known to promote changes in phosphorylation of MAPK pathway and specifically share STE11, STE20 and CDC24 protein kinases (Brückner et al. 2011; Saito & Posas 2012). The metabolic adaptations upon salt perturbation are arguably better characterised than pheromone and these involve the regulation ion membrane transporters and the production and retention of glycerol (Saito & Posas 2012).

Wild-type strains of *S. cerevisiae* displays long periods to initiate the signalling response when stimulated with pheromone, while salt response is almost immediate (Vaga et al. 2014; D’Aquino et al. 2005). To ensure that both responses began at comparable time-scales, yeast strains carrying a CDC28 analog sensitive version were used and CDC28 was inhibited with an ATP analog (Fig 1B). The dynamic response of metabolism was captured for both conditions pairing and expanding the time-points acquired in the phosphoproteomics data-set, i.e. 0 and 25 seconds and 1, 4, 5, 9, 10, 15, 20, 25, 35, and 45 minutes, 0 seconds represents the unperturbed state immediately before the stimuli are added. Cell material was extracted with fast-filtration and analysed with targeted (LC-MS/MS) (Buescher et al. 2010) and untargeted (QTOF-MS) (Fuhrer et al. 2011) mass-spectrometry (see Methods). Robustly identified ions spectra mass were then matched and annotated to an existing database (Fuhrer et al. 2011). In total, we measured with LC-MS/MS 54 metabolites and with QTOF-MS 11,190 ions for which 452 were mapped to metabolites using the genome-scale model iMM904 (Mo et al. 2009). After quality control, we retained 26 metabolites for the downstream analysis from LC-MS/MS and 196 ions mapping to 74 metabolites from the QTOF-MS (see Methods). In order to estimate the reliability of the metabolite measurements, we compared the metabolic fold-changes measured in both targeted and untargeted MS (Fig 1C). A total of 11 unique metabolites were quantified with both methods and these showed strong concordance (spearman’s rho=0.77, *p*-value < 1.9e−44). On the untargeted data-set, 33 ions were defined as significantly changing in at least one of the time-points analyzed (see Methods). These include several examples of metabolites known to be regulated under these conditions (Fig. 1D). In general, there was a lack of measured products and reactants from the same reaction. However, fumarate and malate were reliably measured and both showed similar profiles (Park et al. 2016). Glycerol 3-phosphate displays an accumulation over time under salt stimulation, consistent with known signalling regulation of GPD1 leading to the production of glycerol (Saito & Posas 2012; Kanshin et al. 2015; Mitchell et al. 2015). Yeast cells also produce and accumulate trehalose under different types of stress conditions, including osmotic stress, and this is visible with the trehalose profile (Hohmann 2002; Saito & Posas 2012). While the metabolic implications of the pheromone stimulation in yeast are generally poorly understood, the pheromone MAPK pathway is known to undergo regulation (Merlini et al. 2013; Bardwell 2005). TOR and the pheromone MAPK signalling pathways have been shown to crosstalk (Brückner et al. 2011). Therefore, it is interesting to see that metabolites involved in the biosynthesis of amino-acids, such as, L-glutamine, N-acetyl-L-glutamate and L-citrulline significantly accumulate over time after pheromone stimulation. Some of these have been previously shown to directly influence TOR1 activity (Oliveira, Ludwig, et al. 2015). These results recapitulate previous findings and therefore support the usefulness of this metabolomics data-set to understand the metabolic adaptation to salt and pheromone.

To compare the responses between time-resolved experiments and steady-state genetic perturbations as well as to test the inference methods across different conditions, we expanded the analysis across a range of different cellular perturbations. Salt and pheromone data-sets were integrated with a compendium of biological experiments including time-resolved measurements related to nitrogen metabolism and steady-state genetic perturbations. To this end we considered a panel of 115 K/Ps knockouts, for which molecular changes at the transcript (Kemmeren et al. 2014), phosphorylation (Bodenmiller et al. 2010) and metabolite (Schulz et al. 2014) were characterised (Fig 1A) (see Methods) as well as metabolomics, transcriptomics and phosphoproteomics data-sets for three perturbations around nitrogen metabolism (Oliveira, Ludwig, et al. 2015; Oliveira, Dimopoulos, et al. 2015). In these studies, yeast cells were perturbed by varying the growth medium from poor to rich nitrogen growing conditions (nitrogen upshift) and vice-versa (nitrogen downshift). Yeast cells were also stimulated with Rapamycin, thereby inhibiting TOR1, a condition that resembles the nitrogen downshift (Fig 1A). Combining all the experimental data-sets together, we obtained a total of 143 different conditions for which metabolic, phosphorylation and gene expression measurements are available, except for salt and pheromone conditions where transcriptomics is not available. These data-sets provide the basis for the systematic and comprehensive analysis of transcriptional and post-transcriptional regulatory interactions with metabolism.

### Inferring activity of transcription-factors, kinases and phosphatases

Changes in gene expression and in protein phosphorylation can be combined with metabolic measurements to identify possible regulatory associations. However, identification of functional regulatory interactions is hampered by the fact that expression is a poor proxy for TFs activity (Cheng et al. 2012; Schacht et al. 2014) and phosphorylation sites often display no functional impact in protein activity (Beltrao et al. 2012; Oliveira et al. 2012). Therefore, to circumvent these limitations we have estimated the changes in activity of TFs and K/Ps. Enzymatic activity of K/Ps were estimated resorting to a comprehensive set of manually curated K/Ps-substrates interactions from PhosphoGrid (Sadowski et al. 2013). TF activities were inferred using a regulatory network obtained by combining gene-expression data from TF knock-out experiments and TF binding sites from ChIP-chip experiments (see Methods). The changes in activity of a regulator can be estimated by considering the changes of its targets (Casado et al. 2013; Cheng et al. 2012). For example, by analysing the phosphorylation changes of reported target sites of a protein K/P, one can predict whether the K/P is changing significantly (Fig 2A).

**Figure 2.**
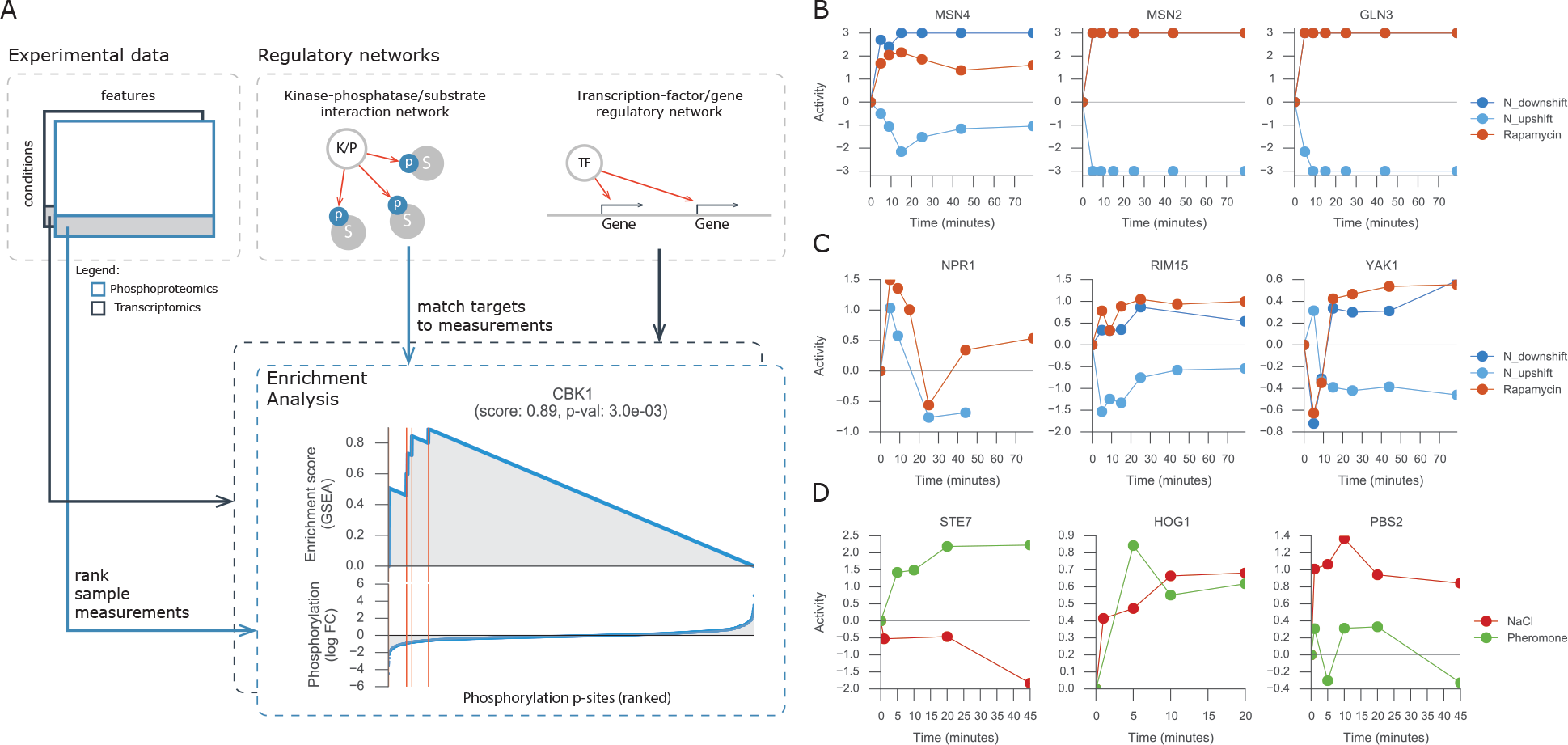
Protein activity analysis. (A) Representation of the workflow used to estimate protein activities using as input an experimental data-set and a regulatory network. Regulatory network will contain either the kinase-phosphatase/substrate interactions or transcription factor/gene associations. GSEA with random permutations is used for each protein in each condition. Red vertical lines represent the targets of the protein in the ascending sorted data-set. (B, C, D) Estimated activity profile of representative proteins for each experiment used in the analysis.

Considering the reported targets of TFs and K/Ps we used the gene-set enrichment analysis (GSEA) (Subramanian et al. 2005) approach to quantify and estimate the significance of the activity of 91 TFs and 103 K/Ps across all conditions (see Methods). The phosphoproteomic data-sets contain 85.2%, 49.2% and 20.0% of missing values in the genetic, nitrogen and salt/pheromone perturbations, respectively. For this reason, the activity scores of K/Ps could not be predicted in 3,227 (48.0%), 498 (25.2%) and 434 (7.7%) cases for the genetic, nitrogen metabolism and salt/pheromone perturbations, respectively. We note however, that the estimated activities do not always rely on measuring the same set of reported targets, and hence the estimated activities matrices are less sparse than the original measurements. For the dynamic experiment of salt and pheromone stimuli there are no transcriptomics available, thus TFs activity could not be calculated.

Nitrogen downshift and rapamycin are similar conditions that inhibit TOR1 activity; in contrast, nitrogen upshift displays increased TOR1 activity. Thus, it is reassuring that the predicted protein activities tend to have similar changes in time for the nitrogen downshift and rapamycin condition, and opposite changes for the nitrogen upshift (Fig 2B, Fig 2C). Several of the predicted activities are in line with known condition dependent activity changes. Examples include the TOR mediated inhibition of MSN2, MSN4 and GLN3 TFs (Beck & Hall 1999) (Fig 2B) and the kinases NPR1 (Schmidt et al. 1998), RIM15 (Pedruzzi et al. 2003) and YAK1 (Martin et al. 2004) (Fig 2C). Moreover, HOG1 and PBS2, central kinases in the response to osmotic stress, display increased activity profiles (Vaga et al. 2014; Saito & Posas 2012) (Fig 2D). Similarly, the STE7 MAPK kinase of the pheromone pathway is predicted to be activated during pheromone stimulation (Fig 2D). These examples suggest that the TFs and K/Ps activities are well predicted and can be used to explore regulatory associations with metabolic changes. Regulator activities provide functional information that can be integrated with metabolic changes to infer functional regulatory interactions. Nevertheless, the activity of the regulators, as their expression and phosphorylation measurements, may be confounded by general cellular states (e.g. growth rate) and therefore lead to indirect associations. In the next section we tested the impact of growth rate on activity estimates and metabolic measurements.

### Growth rate implications in intracellular changes

General effects in the cell, such as cell cycle and growth rate, can act as confounding factors when searching for regulatory associations between TFs and K/Ps and metabolic changes. In particular, gene expression changes, which upon different perturbations have been shown to be tightly correlated with growth rate due to changes in the distribution of cells over the cell cycle phases (O’Duibhir et al. 2014; Brauer et al. 2008). Considering that relative growth rate measurements are available for the genetic perturbations experiments and for each time point of the dynamic nitrogen metabolism experiments, we set out to assess how much of the variation in the data-sets can be explained by growth rate alone. To this end, we performed Principal Component Analysis (PCA) on TF and K/P activities as well as metabolomics measurements. We then measured the correlation between relative growth rate and each of the top three principal components (PCs) (Fig S1). Genetic perturbations metabolomics data-set PC 1 displayed a moderate correlation (Pearson’s r=0.25, *p*-value < 7.0−e3) with the relative growth rates of the knockout strains. Growth rate displayed stronger correlations (Pearson’s r=0.35 and −0.54, *p*-values < 1.2e−4 and 1.1e−4) with PC 1 of K/P and TF activities (Fig S1). The same analysis was performed for the dynamic nitrogen metabolism data-set where metabolomics PC 2 displayed a strong correlation with the relative growth rate over time (Pearson’s r=0.72, *p*-value < 8.0e−4). For the estimated K/P and TF activities, PC3 and PC2 showed also strong correlations with growth (Pearson’s r of 0.51 and 0.69, *p*-values < 3.2e−2 and 1.6e−3) (Fig S1).

In summary, the variance in molecular measurements for the steady-state genetic perturbation experiments is more strongly influenced by the growth rate than the measurements performed in the dynamical perturbations. For the steady-state conditions, the gene-expression changes are the molecular changes, confounded mostly by the growth rate.

For the subsequent association analyses we tested the impact of removing the growth rate from each data-set to rule out any confounding effects it may have on the identification of direct functional interactions. To this end, we regressed-out growth rates from the original metabolite measurements and estimated TFs and K/Ps activities using linear regression models and growth as a covariate.

### Estimating metabolic changes from transcription-factors, kinases and phosphatases activities

Next we explored the correlations between TFs and K/Ps enzymatic activities and metabolic changes for each of the three experiments: genetic, nitrogen metabolism and salt/pheromone perturbations. To identify the relationships we used linear regression models that consider the estimated activities as features and metabolite fold-changes as observations. Considering the low number of samples available, specifically for the time resolved experiments, a crossvalidation procedure of leave-one-out (LOO) was used (see Methods). This allowed us to understand how much information can be transferred within each experiment to predict the metabolite variations in an independent testing sample. Thus, for each experiment and each metabolite, independent training and test data-sets were generated leaving one sample out at a time for test, i.e. single KO or time-point, and thereby generating a complete metabolomics matrix with estimated fold-change values (Fig 3A). The analysis was performed using TFs and K/Ps activities independently. In each experiment four different types of input matrices are used to predict each metabolite, i.e. K/P or TF activities with and without growth normalisation, with the exception of the dynamic experiment with NaCl and pheromone for which neither growth rate nor transcriptomics measurements were available. To minimize possible effects of over fitting while training the linear models an Elastic Net feature regularisation approach was used (Methods).

**Figure 3.**
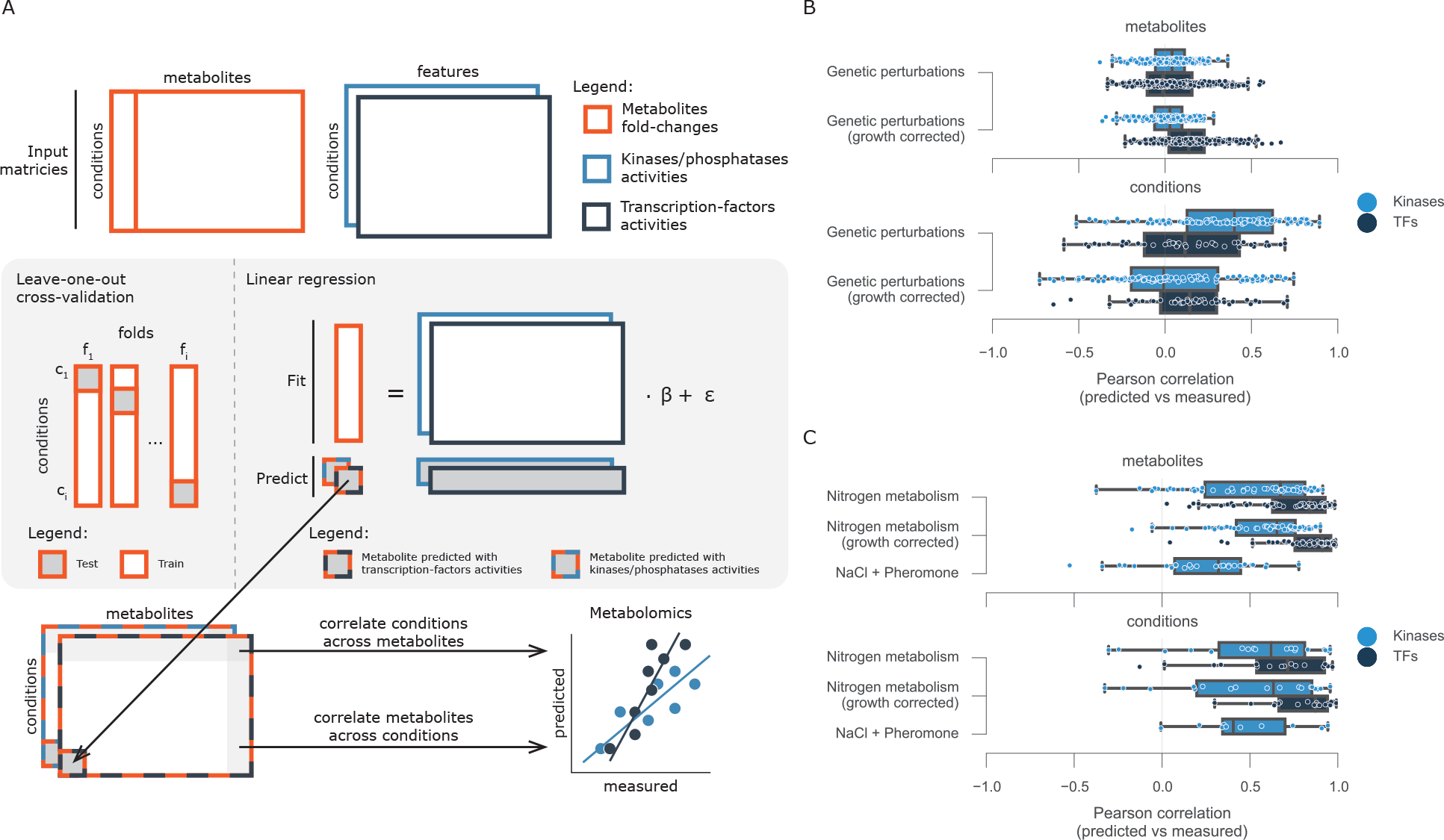
Computational framework for finding associations between protein and metabolites. (A) Diagram of the analysis. For each metabolite measured a multilinear regression analysis was performed using leave-one-out cross-validation. Changes in activity for kinases/phosphatases and transcription factors were used independently to estimate the metabolite fold-change. Independently predicted metabolite fold-change matrices were then correlated metabolite and condition wise with the measured values. (B) Distributions of the correlation values between predicted and measured metabolite fold-changes.

Firstly, we considered the genetic perturbations and assessed the capacity of TFs and K/Ps activities to predict the changes of a given metabolite across the panel of knockouts (Fig 3B, metabolites). This would be, for example, changes in concentration of glutamine across the knockout conditions. We evaluated the capacity of the models by correlating the independently predicted fold-changes to the observed ones. This procedure was performed for each metabolite and the results were summarized as correlation distributions. The metabolite variation across the different conditions were generally poorly predicted using either the TFs or K/P activities, displaying median correlations close to zero (Fig 3B). This did not change when we used the growth corrected data. We then measured how well the models predict the changes in all metabolites in a given condition (Fig 3B, conditions). This tests the capacity to, for example, predict the changes of all metabolite changes in HOG1 knockout. Overall, we obtained similar results as with the metabolites analysis, one difference is the improvement in predictive power considering K/Ps activities. However, this increase is mostly lost when we use the growth corrected data.

A similar analysis as with genetic perturbations was applied to the dynamic experiments to estimate the metabolic variation. The trained models displayed in general higher predictive power than the genetic perturbations (Fig 3C). Overall, in the dynamic nitrogen experiments, TFs displayed better agreement between measured and predicted metabolite fold-changes than K/Ps, across metabolites (Fig 3C, metabolites) and across conditions (Fig 3C, conditions). Also, models trained with growth normalised activities obtained similar results to non-normalised datasets. The metabolic changes in the salt and pheromone experiment could be reasonably explained using the K/P activities across metabolites (Pearson’s r=0.32) (Fig 3C, metabolites) and conditions (Pearson’s r=0.41) (Fig 3C, metabolites). These were generally worse than the nitrogen experiment, and could be a consequence of the lower number of significantly changing metabolites when compared to the nitrogen experiments.

The predictive difference between growth normalised and non-normalised K/Ps activities in the genetic perturbations (Fig 3B, conditions) suggest that associations important to predict a new condition are generally dependent on global growth effects, and thereby likely to be indirect. Furthermore, the different predictive power between TFs and K/Ps on the dynamic nitrogen experiments suggest that changes in TF activities are more predictive of metabolic changes. Nevertheless, one needs to consider that very different technologies are used to measure the underlying data-sets, i.e. transcriptomics and phosphoproteomics and this may impact the predictive power of the data-sets.

### Inferring putative regulatory protein-metabolites interactions

Considering that the metabolic predictions based on time-resolved experiments partially circumvented indirect effects and displayed the best predictive power (Fig 3B, Fig 3C) we decided to focus on these data-sets for the inference of protein-metabolite regulatory interactions. Moreover, since growth has been shown to possibly act as a confounding effect (Fig 3B) we only used the data-set with growth normalised for the nitrogen metabolism experiments. We also considered TFs and K/Ps separately and searched for putative regulatory associations with the metabolite changes.

We started by investigating the capacity of the TFs activities to estimate the metabolites fold changes in each nitrogen related perturbation. To this end, we used a learning procedure, analogous to the one used before, but instead of LOO, a three-fold cross-validation was used to leave each of the environmental perturbations out at a time (see Methods). This was performed independently for each metabolite and the agreement between the measured and predicted values was calculated using Pearson correlation coefficients (Fig 4A). Consistently with the previous analysis, a large fraction of the metabolites were well predicted in downshift and rapamycin conditions. The best performances are obtained in the nitrogen downshift and the rapamycin experiments with similar median correlations. This could be expected since these are related conditions and the relationships learned from one may more readily apply to the other. Then, we considered only the best predicted metabolites (Fig S2A) and explored putative protein-metabolite associations using all the three nitrogen conditions together with bootstrapped linear regression models (see Methods). The associations were estimated 20 times with 80% of the samples randomly selected, therefore generating 20 coefficients for each TF-metabolite association. The average of the TF-metabolite coefficients represents a confidence score on the association (Fig 4B).

From the reported associations, LEU3 involved in the biosynthesis of leucine is positively associated with several metabolites involved in the biosynthesis of amino-acids, e.g. L-glutamine, L-citrulline, ornithine (Friden & Schimmel 1988; Nielsen et al. 2001). Also, the involvement of PUT3 in the proline utilisation pathways and its positive association with L-proline is captured by the linear model coefficients (Huang & Brandriss 2000; Axelrod et al. 1991; Siddiqui & Brandriss 1989). These results seem to confirm that the regulatory interactions found are biologically relevant, although they can be a result of direct or indirect associations. For example, a direct interaction can occur if a TF regulates the expression of metabolic enzymes and thereby controls directly metabolite concentration. The association can occur in the opposite direction where metabolites can directly regulate the activity of TFs. In contrast, indirect associations can be established, for instance, if metabolite changes are a consequence of downstream effects of TFs or if a cell state results in changes of both TF activity and metabolite concentration independently. In order to study this, we firstly identified the enzymes that use or produce each measured metabolite and considered a list of known TF-target proteins (from our assembled TF regulatory network), TF-gene genetic interactions (from BioGRID (Chatr-Aryamontri et al. 2015; Oughtred, Chatr-aryamontri, et al. 2016; Oughtred, Chatr-Aryamontri, et al. 2016)) or TF-gene functional interactions (from STRING (Jensen et al. 2009)) (see Methods). We then searched for enrichment of known TF-target, TF-gene genetic and functional associations among the top predicted TF-enzyme-metabolite interactions (Fig S3A). No significant association was found. We also note that the variation in TFs activities are almost fully explained by the first PC that captures 85.6% of the total variance in the data (Fig S1). Furthermore, TFs activities showed similar profiles within TOR1 inhibition conditions, nitrogen downshift and rapamycin, and opposing profiles in TOR1 activation condition, nitrogen upshift. Hence, this shows lack of specificity in the gene expression response and can partially explain the limited capacity to identify direct associations with metabolites. However, regressing-out the first principal component from the TFs activity scores and from the metabolomics measurements did not improve the enrichment in direct TF-target associations (data not shown). These findings support the idea that although the TFs activities are predictive of metabolic changes these relationships are likely to be indirectly due to changes in cellular states or via transcriptional regulation of genes that are not those immediately in the vicinity of the associated metabolites.

For the K/P-metabolite associations we used all five dynamic perturbations: nitrogen upshift, nitrogen downshift, rapamycin, NaCl and pheromone (Fig 4C). With the exception of the pheromone the other conditions showed similar median correlations between the measured and predicted metabolites fold-changes. However, the performance is overall lower than for the leave-one-out test (Fig 3C), as would be expected from a more stringent evaluation. The top predicted metabolites were selected (Fig S2B) and an analogous approach used for the TFs was used to identify K/P-metabolite associations (see Methods) (Fig 4D).

RIM15 and TPK1 displayed the strongest associations with the metabolites and these play a key role in the regulation of the cellular growth and their adaptation to nutrient availability (Chavel et al. 2014; Conrad et al. 2014; Broach 2012). TPK1 inhibits the activity of RIM15 to regulate cell cycle, thus this justifies that both display opposite associations (Pearson’s r −0.92, *p*-value < 8.6e−6). Furthermore, RIM15 is inhibited by TOR1 (Swinnen et al. 2006; Broach 2012) and considering that L-proline is a poor nitrogen source leading to decreased TOR1 activity, this is consistent with the positive association between RIM15 and L-proline, and that TPK1 displays the inverse. Of note, TOR1 and RIM15 display similar metabolite relationships despite their inverse biological association, this happens because TOR1 activity is wrongly estimated due to lack of robustly measured targets. This is emphasised with the non-significant negative correlation between TOR1 and RIM15 activity scores (Pearson’s r −0.17, *p*-value < 3.98e−1). The associations may be direct causal K/P-enzyme-metabolite relationships, but they could also be indirect or in the opposite direction where a metabolite change impact kinase activity. We performed an enrichment analysis similar to the one described before, but now considering from BioGRID genetic and physical interactions. For each K/P-gene network we tested for enrichment of true interactions in the top-predicted K/P-enzyme-metabolite associations (Fig S3B). We observed a significant but weak enrichment for functional interactions (AROC=0.63, Fig S3B) and direct K/P-target relationships (AROC=0.61, Fig S3B). This significant enrichment in known K/P-target associations contrasts to the inferred TF-metabolites associations above. Specifically, from 16 functional interactions reported in STRING overlapping in our set of inferred associations half of those displayed a positive absolute coefficient. These results suggest that the retrieved associations contain some direct K/P-target relationships.

TFs-metabolites and K/Ps-metabolites associations can also be taken together to elucidate associations between K/Ps and TFs. For example, TPK1 kinase inhibits the activity of TF ADR1 (Conrad et al. 2014) and these have inverse associations considering the five top predicted metabolites that they share (Pearson’s r −0.92, *p*-value < 2.6e−2). Also, YAK1 is required for full activation of MSN2/4 TFs (Broach 2012) and this association is visible, although not significant, between MSN4 and YAK1 metabolites associations (Pearson’s r 0.8, *p*-value < 1.1e−1), for example, both have negative associations with dUDP and positive associations with L-Proline. However, this type of analysis is limited to the number of metabolites that can be well predicted with the TFs and K/Ps activities.

In summary, the estimated activity of TFs and K/Ps were capable of building predictive models to infer the metabolic adaptations of previously unseen conditions across different dynamic experiments. Specifically, TFs activities provided better predictive power than K/Ps, although K/Ps-metabolite associations are more likely to represent previously reported interactions.

## Discussion

Signal transduction is an important cellular mechanism that allows cells to sense and respond to environmental cues. These mediate intracellular adaptations by regulating a variety of biological processes, including, metabolism and gene expression. Thereby interactions among different biological processes occur and are very important to coordinate the whole phenotype of the cell. Nevertheless, the systematic identification and functional annotation of these regulatory interactions is still a challenge. Experimental data-sets covering different omics in similar conditions are becoming more available and it is likely that analyses like the one we propose here will be useful to systematically explore these regulatory events.

The key novelty of the approach proposed here is that regulatory interactions are inferred from estimated activity of TFs and K/Ps, which are difficult to measure directly. This provides the possibility of considering the activity profile of regulatory proteins important for the experimental conditions at hand which has not been considered in previous studies of metabolism using phosphoproteomics and transcriptomics.

Our results show that it is possible to use K/Ps and TFs activities to predict changes of several metabolites in time-resolved experiments. However, the predictive power does not extend to all conditions. For example, the models trained with K/Ps activities showed limited capacity in the pheromone perturbation experiment. This can arise from the higher technical variability in the data obtained and also be due to lower number of regulated metabolites. Additionally, the regulatory interactions are often condition specific. As such, if the proteins that are important to regulate the nitrogen or osmotic related conditions are not used for the pheromone response, then the associations learned cannot be predictive of the pheromone induced metabolic changes. Interestingly, protein-metabolite interactions inferred from the genetic perturbations experiment displayed poor predictive power to estimate the metabolic changes of a new condition, in contrast to the dynamic experiments (Fig 3B, Fig 3C). This suggests that time-resolved experiments provide a more efficient design to infer regulatory associations by circumventing general confounding effects that can be seen in the steady-state.

Nevertheless, while the protein-metabolite interactions that we infer provide reasonable power to predict metabolic changes (Fig 4A, Fig 4C) of unseen conditions the predicted regulator-enzyme-metabolite interactions are not strongly enriched in previously regulatory interactions. Some features, particularly kinases or phosphatases activities, such as RIM15 and TPK1, were important features to estimate metabolites fold-change. This reassuringly assesses that the estimated protein activities profiles are biologically relevant and useful for inferring metabolic adaptation in novel conditions. The time-resolved metabolomics experiment under salt and pheromone resulted in only moderate metabolic changes, when compared to the nitrogen conditions. This smaller variation may explain the lower power in identifying regulator-metabolite associations in these conditions. We believe that this emphasises the importance of designing experiments that adequately perturb both signalling and metabolism, without possible confounding effects, such as CDC28 inhibition. Another possible limitation of this approach is that, while we used comprehensive resources, we only considered prior knowledge of reported K/Ps-substrate and TFs-gene regulatory interactions. Furthermore, the lack of missing values in transcriptomics data-sets provides increased robustness to TFs activities when compared to K/Ps activities. Nevertheless, both protein activity profiles showed comparable predictive power, thus partially guaranteeing that the existing bias does not penalise greatly the K/Ps.

**Figure 4.**
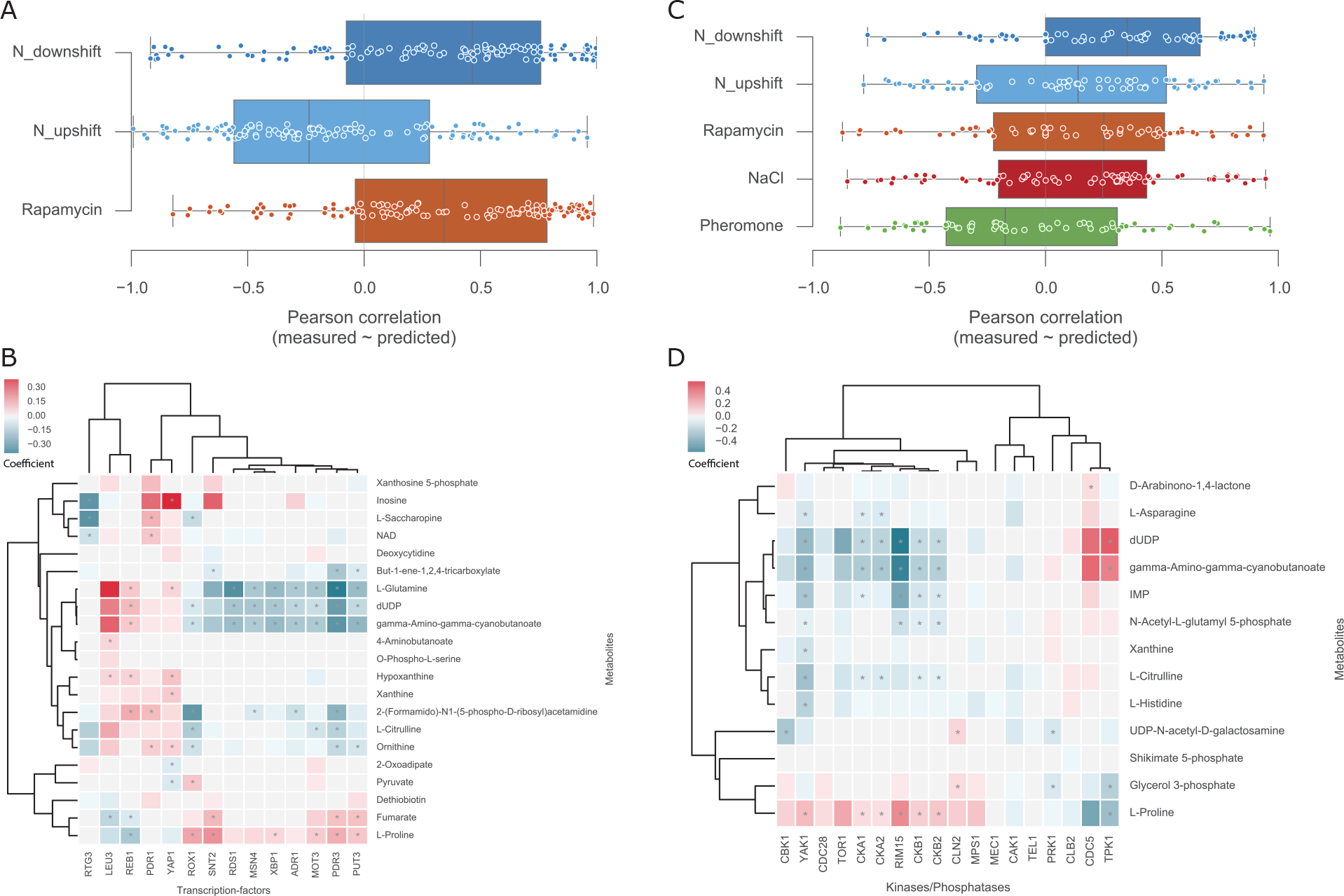
Overview of the putative protein-metabolite regulatory interactions. (A) Distribution of the metabolite predicted and measured correlations using 3-fold cross-validation leaving each condition out at a time using the TFs activities scores. (B) Heatmap of the TFs-metabolites associations where values represent the averaged coefficients. (C) Correlation distributions between predicted and measured using K/Ps activities with 5-fold cross-validation leaving each condition out at a time. (D) Heatmap of the K/Ps-metabolites associations where values represent the averaged coefficients. Coefficients distributions are calculated using a bootstrap cross-validation randomly leaving 20% of all the samples out. This procedure is performed twenty times and the coefficients are then averaged. Asterisks (*) identify significant, FDR < 5%, Pearson correlations between the activity profiles and the metabolite fold-change across all conditions.

The increasing number of recorded interactions for regulators will provide important information to expand the coverage of TFs and K/Ps for which it is possible to estimate activities and increase the robustness of the estimated activity. The generic characteristic of the approach used to infer regulatory interactions allows it to be easily expanded to integrate other types of information and thereby augment its predictive power to infer causal and direct protein-metabolite interactions. For example, to account for other confounding effects, such as, cell cycle as a covariate and thereby remove it as a possible source of interactions. Furthermore, other types of biological measurements can also be integrated, for example, protein abundance. This can provide information into other regulatory mechanisms but also provide information to possible associations between the different regulatory processes, for instance, phosphoproteomics measurements are intrinsically dependent on protein abundance.

In this study we demonstrated the utility of phosphoproteomics and transcriptomics data-sets to estimate the enzymatic activity of K/Ps and TFs, respectively. The estimated activities recapitulated several previously expected regulatory events, such as, HOG1 and PBS2 responses to osmotic stress, and RIM15 and MSN2/4 activation under TOR1 inhibition. This results emphasise the usefulness of this approach to explore functional implications in regulatory proteins. Our results also showed that activity profiles are informative features to estimate metabolite changes in dynamic experiments. Interestingly, the same was not visible across a large panel of K/Ps knockouts, supporting the idea that time-resolved experiments are a better experimental design for the identification of causal regulatory interactions. We expanded on previous work by developing a novel and rigorous framework to identify regulatory associations between the estimated activities and metabolite changes. Rigorous analysis of the putative TFs-metabolite and K/Ps-metabolite associations occurring in the dynamic experiments revealed that despite their regulatory implication these are most likely indirect. Further confirmation of these results with the integration of other experimental data-sets will provide deeper insights into the regulatory events mediating the metabolic phenotype.

## Methods

### Strain, growth and sample preparation

The *Saccharomyces cerevisiae* strain used for the salt and pheromone dynamic experiments was BY4741 as in Vaga et al. (Vaga et al. 2014; D’Aquino et al. 2005), this strain is provided with a CDC28-as allele that can be directly inhibited by means of 1-NA-PP1, the ATP analog “PP1 analog 8”. Cells were grown in 500-ml shake flasks at 30°C in 50 ml SD medium to an 0D600 of 0.6. The ATP analog was added to a final concentration of 10μM. One hour after CDC28 inhibition cells were perturbed with NaCl to a final concentration of 0.4M or pheromone to a final concentration of 1 μM. Cells were extracted by vacuum-filtering culture aliquots on a 0.45 μm pore size nitrocellulose filter (Millipore). The filter was immediately transferred to 3 ml 2:2:1 MeOH/AcN/ddH2O precooled at −30°C. Samples for LC-MS/MS were supplemented with 200 pl uniformly-labeled 13C *E. coli* extract as internal standard and dried completely in a vacuum centrifuge (Christ-RVC 2-33 CD plus, Kuehner AG, Birsfelden, Switzerland). The dried extracts were resuspended in 100 μl MilliQ water before analysis.

### Acquisition of intracellular metabolite levels

Targeted metabolomics was performed by LC-MS/MS as described before (Buescher et al. 2010). The mass-spectrometer was operated in negative mode. Data acquisition and peak integration were performed with the Xcalibur software version 2.07 SP1 (Thermo Fisher Scientific) and in-house integration software. Metabolite peak areas were normalized to uniformly-labeled 13C internal standards.

Untargeted metabolomics was performed by direct flow double injection of extracts on an Agilent 6550 series quadrupole TOF MS operated in negative mode. Detected ions were annotated against the yeast metabolic reconstruction iMM904 (Mo et al. 2009) with a stringent tolerance of 0.001 Da. Considering only highly confidently detected ions for annotation, 196 ions were annotated to 270 yeast metabolites (Supplementary Table 1). The overall performance of the untargeted metabolomics was compared to the targeted metabolomics (Fig 1D). High concentrations of salt in the NaCl perturbation experiment resulted in a strong effect on the ion matrix in the QTOF-MS measurements. To prevent this matrix effect from affecting data analysis we normalised the data to the second time-point (25 seconds) instead of the 0 seconds timepoint.

Statistical significance of the ion fold-changes for the QTOF-MS measurements was estimated with a two-sided t-test followed by multiple hypothesis correction with false-discovery rate. The list was then filtered and only ions with and an absolute fold-change higher than 1 were considered, resulting in 33 ions.

### Compendium of yeast data-sets

For K/Ps knockouts in yeast, a total of 3,011 transcripts were measured across 1,484 deletion mutants, comprising approximately 26.4% of all protein-coding genes in yeast (Byrne & Wolfe 2005). Phosphoproteomics profiles of 125 K/P knockouts, as compared to a wild-type strain, were acquired using label free mass-spectrometry (LC-MS/MS) measuring 4,263 unique single phosphorylated phosphosites in at least one condition. Intracellular measurements of metabolites were obtained during the exponential growth phase and analyzed using nontargeted direct injection and time-of-flight mass-spectrometry (QTOF-MS). In total, 1,698 unique ions were detected across 118 kinases/phosphatases knockouts. Metabolomics and phosphoproteomics data-sets overlap in 115 knock-out conditions, and metabolomics intersects the transcriptomics in 45 knock-out conditions.

Dynamic perturbations to nitrogen metabolism and TOR signalling were captured in a time frame from 0 to 79 minutes, where 0 minutes represents the unperturbed state. Transcriptomics measurements covered 5,620 transcripts across all the time-points of the conditions. Phosphoproteomics captured the profile of 1,660 single phosphorylated phosphosites (84.8% serines, 14.2% threonines and 1.0% tyrosines) over the same time-points. Given the lack of complete coverage, only 50.8% of the whole matrix is measured. Intracellular metabolomics were acquired with QTOF-MS and quantified a total of 146 ions, after quality filtering, across all conditions and time-points.

### Activity inference method

Kinases/phosphatases and transcription factor activities were estimated using GSEA approach (Subramanian et al. 2005) and statistical significance was calculated against a null hypothesis generated by randomising 1000 times the regulator reported targets (Subramanian et al. 2005). Activity scores were calculated using the log10 of the empirical *p*-value and signed according to the direction of the enrichment, for example, regulators enriched towards negative fold-changes have a negative score. K/Ps target phosphosites were extracted from PhosphoGrid (Sadowski et al. 2013). Specificities for a total of 177 transcription factors were collected in form of a position weight matrices (PWMs) from JASPAR (Mathelier et al. 2014). Weight matrices were trimmed to remove consecutive stretches of low information content (<0.2) on either end. The log-scoring scheme defined in (Wasserman & Sandelin 2004), was used to score potential target sequences against weight matrices. The log score is normalised to the best and worst matching sequence to the weight matrix, resulting in a value that lies between 0 and 1, where 1 denotes strong binding to the matrix and 0 denotes no binding. Genome wide gene expression profiles for 837 gene-knockout strains were collected from three studies (Kemmeren et al. 2014; Hu et al. 2007; Chua et al. 2006), of which 148/837 were a known transcription factor with a defined specificity weight matrix. Studies provided either a Z-score or *p*-value for each gene as a measure of over or under-expression, relative to the distribution of values for all genes. Two-tailed *p*-values were computed from Z-scores when a p-value was not provided (Chua et al. 2006). In cases where TF knockout was repeated between studies, the lowest p-value for each gene was used. ChIP-ChIP tracks for 355 proteins were collected from four studies (Harbison et al. 2004; Rhee & Pugh 2011; Tachibana et al. 2005; Venters et al. 2011), via the Saccharomyces genome database (Christie et al. 2004). 144/355 of proteins were transcription factors with a defined specificity weight matrix. The TF-gene network was then defined as all TF-gene pairs with a p-value below 0.01 and contained a ChIP-ChIP region upstream of the regulated gene, which scored highly against the weight matrix of the TF (normalised logscore>0.9).

### Linear regression methods for estimating metabolic changes

Python module Sklearn (Pedregosa et al. 2011) version 0.16.1 was used to perform linear regression analysis and default parameters were used unless stated otherwise. Linear models with combined L1 and L2 regularization, Elastic Net, was used with the |1_ratio of 0.5. Elastic net regularization simplifies the complexity of the model by removing the least important features, similar to Lasso regularisation, but also considering a L2 regularization, similar to Ridge, to avoid random feature elimination when collinearity exists among the features.

To infer the predictive power within each data-set across all the measured ions (Fig 3B), for each metabolomics data-set, ions displaying low variation across the samples were discarded by considering only those that showed a standard deviation higher than 0.4. KPs activities were filtered to only consider kinases or phosphatases with an activity score estimated in at least 75% of the samples of each data-set, the remaining missing values were replaced with zeros for the machine learning approaches. For the Elastic net regressions different alphas were tested and an alpha of 0.01 obtained the best overall performance, therefore this was used in all the models. For each metabolite a leave-one-out cross-validation was used, thus all but one sample were used to train the linear regression model and then the test sample was used to estimate the metabolite fold-change. Performing this systematically across all metabolites and conditions generated a predicted matrix for which each value is estimated independently. The agreement between the measured and predicted ions fold-changes was calculated with Pearson correlation coefficients across rows (ions) and columns (conditions).

### Linear regression models to predict K/Ps-metabolites and TFs-metabolites associations

Protein-metabolite associations were inferred only using the time-resolved metabolomics datasets. For the TFs activities only the nitrogen metabolism perturbations were used considering that no transcriptomics data was available for the NaCl/Pheromone perturbations. For this analysis no filtering was applied in the metabolomics data-sets and K/Ps activities were filtered as before to consider only those consistently measured and estimated across 75% of the conditions. The capacity of predicting each ion fold-change in each condition was tested using k-fold cross-validation, where each fold corresponds to the time-points measured in each condition, thus 3 and 5 folds were used for the TFs and KPs activities, respectively. Elastic net models were used and the alpha was estimated using a bootstrap approach of ten iterations leaving out 20% of the samples. A range of 100 alphas was considered as default by the Sklearn python module (Pedregosa et al. 2011). Train and test features, TFs and KPs activities, were standardised, and the observed variables, metabolomics, were centered before training the linear model. The predictive power of each ion in each condition was estimated by using the k-fold models, inferring the agreement between predicted and measured in the left-out condition using Pearson coefficient and coefficient of determination metrics. The top predicted ions were those that displayed a Pearson correlation *p*-value lower than 0.05 and an coefficient of determination higher than zero.

For the top predicted ions the feature importance was estimated using all the conditions together with two bootstraps. The first bootstrap was 20 iterations and leaves out 20% of the samples out, for each iteration an inner bootstrap with 10 iterations leaving out another 20% of the data is performed to estimate the alpha of the Elastic net. This estimates 20 coefficients for each feature-metabolite association. As before, train features and observations are standardised and centered. The most important features per ion are estimated by taking the median of the coefficients and the Pearson correlation between the protein activity and the ion fold-change.

### Data and analysis code availability

All data analysis was performed in Python (v 2.7.10) and all the code, preprocessed data-sets and generated plots are openly available in github under the GNU general public license version 3 in the following URL https://github.com/saezlab/yeastphospho. All plotting was performed using Python modules Matplotlib version 1.4.3 (Hunter 2007) and Seaborn version 0.7.0 (Waskom et al. 2014).

## Acknowledgements

This work was supported by an EMBO Short Term Fellowship (EMBO ASTF 285-2015) and by EMBL PhD programme.

### Author contributions

EG performed the computational analysis. EG and ZR performed the biological experiments. ZR, MZ and US provided experimental supervision. OW assembled the transcriptional regulatory network. DO, PB and JSR supervised the computational analyses. EG, PB and JSR wrote the manuscript. All authors read and contributed to the manuscript.

---

**Supplementary figure 1.**
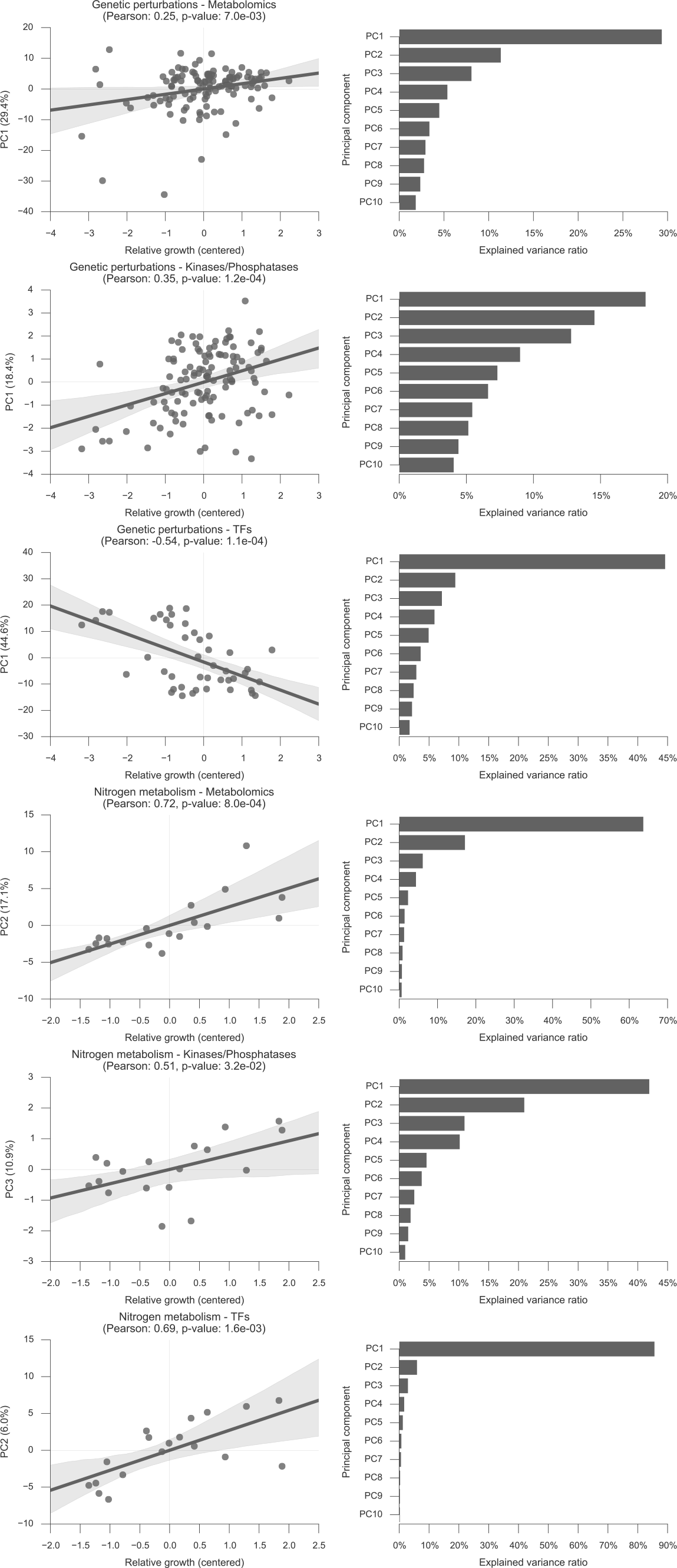
Principal component analysis of of data-sets and correlation with relative growth rate. The principal component with higher absolute correlation coefficient was picked and plotted.

**Supplementary figure 2.**
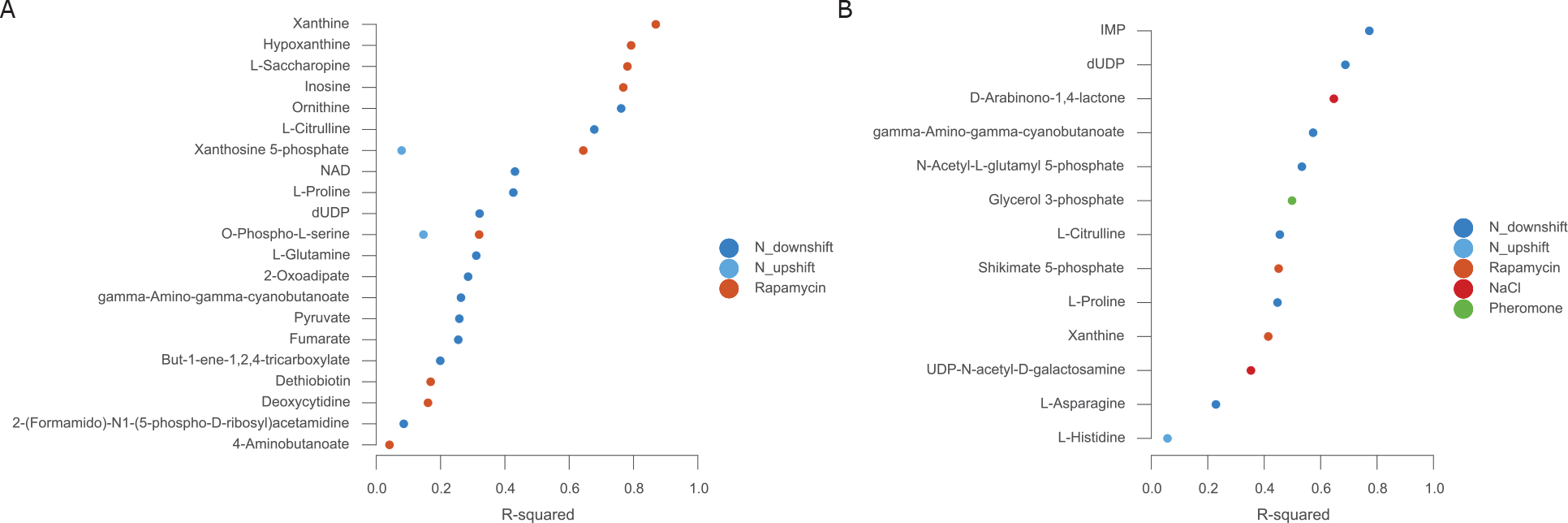
List of top predicted metabolites using a) TFs activities and b) K/Ps activities. List of metabolites that displayed a positive coefficient of determination and significant Pearson correlation between the measured and predicted fold-changes across the different conditions.

**Supplementary figure 3.**
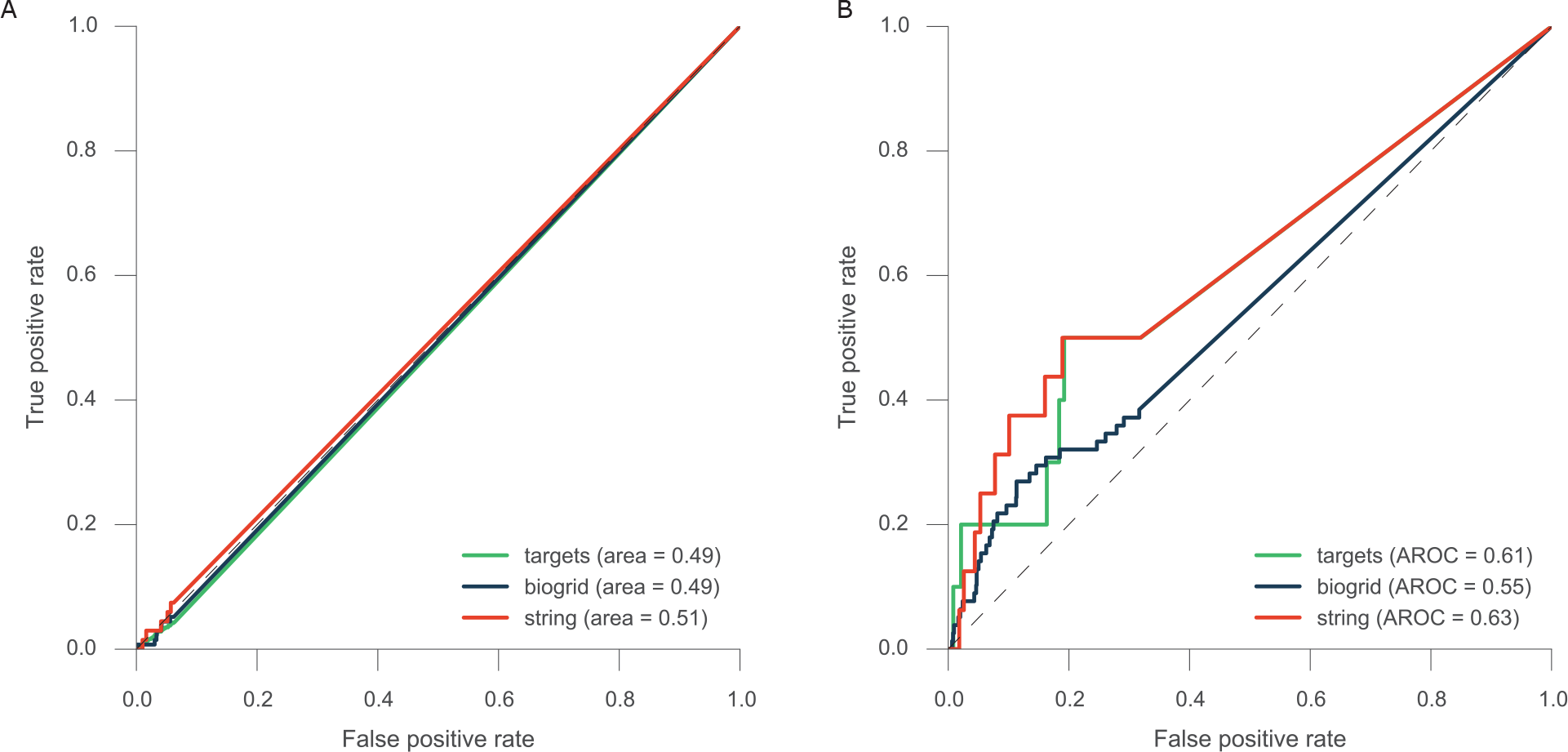
ROC-curve analysis of the average feature coefficients. True-positive tables were built considering the specified resources.

## Tables and their legends

Table 1. Salt and pheromone metabolomics experiments in yeast and metabolites annotation.

Table 2. Kinases/phosphatases and transcription factor activity scores.

Table 3. Protein-metabolites interactions betas.

## References

Axelrod J.D., Majors J. & Brandriss M.C., 1991. Proline-independent binding of PUT3 transcriptional activator protein detected by footprinting in vivo. Molecular and cellular biology, 11(1), pp.564–567.

Bardwell L., 2005. A walk-through of the yeast mating pheromone response pathway. Peptides, 26(2), pp.339–350.

Beck T. & Hall M.N., 1999. The TOR signalling pathway controls nuclear localization of nutrient-regulated transcription factors. Nature, 402(6762), pp.689–692.

Beltrao P. et al., 2012. Systematic functional prioritization of protein posttranslational modifications. Cell, 150(2), pp.413–425.

Bodenmiller B. et al., 2010. Phosphoproteomic Analysis Reveals Interconnected System-Wide Responses to Perturbations of Kinases and Phosphatases in Yeast. Science signaling, 3(153), pp.rs4–rs4.

Brauer M.J. et al., 2008. Coordination of growth rate, cell cycle, stress response, and metabolic activity in yeast. Molecular biology of the cell, 19(1), pp.352–367.

Broach J.R., 2012. Nutritional control of growth and development in yeast. Genetics, 192(1), pp.73–105.

Brückner S. et al., 2011. The TEA transcription factor Tec1 links TOR and MAPK pathways to coordinate yeast development. Genetics, 189(2), pp.479–494.

Buescher J.M. et al., 2010. Ultrahigh performance liquid chromatography-tandem mass spectrometry method for fast and robust quantification of anionic and aromatic metabolites. Analytical chemistry, 82(11), pp.4403–4412.

Byrne K.P. & Wolfe K.H., 2005. The Yeast Gene Order Browser: combining curated homology and syntenic context reveals gene fate in polyploid species. Genome research, 15(10), pp.1456–1461.

Casado P. et al., 2013. Kinase-substrate enrichment analysis provides insights into the heterogeneity of signaling pathway activation in leukemia cells. Science signaling, 6(268), p.rs6.

Chatr-Aryamontri A. et al., 2015. The BioGRID interaction database: 2015 update. Nucleic acids research, 43(Database issue), pp.D470–8.

Chavel C.A. et al., 2014. Global regulation of a differentiation MAPK pathway in yeast. Genetics, 198(3), pp.1309–1328.

Cheng C. et al., 2012. Understanding transcriptional regulation by integrative analysis of transcription factor binding data. Genome research, 22(9), pp.1658–1667.

Christie K.R. et al., 2004. Saccharomyces Genome Database (SGD) provides tools to identify and analyze sequences from Saccharomyces cerevisiae and related sequences from other organisms. Nucleic acids research, 32(Database issue), pp.D311–4.

Chua G. et al., 2006. Identifying transcription factor functions and targets by phenotypic activation. Proceedings of the National Academy of Sciences of the United States of America, 103(32), pp.12045–12050.

Chubukov V. et al., 2014. Coordination of microbial metabolism. Nature reviews. Microbiology, 12(5), pp.327–340.

Conrad M. et al., 2014. Nutrient sensing and signaling in the yeast Saccharomyces cerevisiae. FEMS microbiology reviews, 38(2), pp.254–299.

D’Aquino K.E. et al., 2005. The Protein Kinase Kin4 Inhibits Exit from Mitosis in Response to Spindle Position Defects. Molecular cell, 19(2), pp.223–234.

Daran-Lapujade P. et al., 2004. Role of transcriptional regulation in controlling fluxes in central carbon metabolism of Saccharomyces cerevisiae. A chemostat culture study. The Journal of biological chemistry, 279(10), pp.9125–9138.

Daran-Lapujade P. et al., 2007. The fluxes through glycolytic enzymes in Saccharomyces cerevisiae are predominantly regulated at posttranscriptional levels. Proceedings of the National Academy of Sciences of the United States of America, 104(40), pp.15753–15758.

Friden P. & Schimmel P., 1988. LEU3 of Saccharomyces cerevisiae activates multiple genes for branched-chain amino acid biosynthesis by binding to a common decanucleotide core sequence. Molecular and cellular biology, 8(7), pp.2690–2697.

Fuhrer T. et al., 2011. High-Throughput, Accurate Mass Metabolome Profiling of Cellular Extracts by Flow Injection-Time-of-Flight Mass Spectrometry. Analytical chemistry, 83(18), pp.7074–7080.

Gerosa L. et al., 2015. Pseudo-transition Analysis Identifies the Key Regulators of DynamicMetabolic Adaptations from Steady-State Data. Cell Systems, 1(4), pp.270–282.

Harbison C.T. et al., 2004. Transcriptional regulatory code of a eukaryotic genome. Nature, 431(7004), pp.99–104.

Herrgård M.J. et al., 2008. A consensus yeast metabolic network reconstruction obtained from a community approach to systems biology. Nature biotechnology, 26(10), pp.1155–1160.

Hohmann S., 2002. Osmotic stress signaling and osmoadaptation in yeasts. Microbiology and molecular biology reviews: MMBR, 66(2), pp.300–372.

Huang H.L. & Brandriss M.C., 2000. The regulator of the yeast proline utilization pathway is differentially phosphorylated in response to the quality of the nitrogen source. Molecular and cellular biology, 20(3), pp.892–899.

Hunter J.D., 2007. Matplotlib: A 2D Graphics Environment. Computing in science & engineering, 9(3), pp.90–95.

Hu Z., Killion P.J. & Iyer V.R., 2007. Genetic reconstruction of a functional transcriptional regulatory network. Nature genetics, 39(5), pp.683–687.

Jensen L.J. et al., 2009. STRING 8--a global view on proteins and their functional interactions in 630 organisms. Nucleic acids research, 37(Database issue), pp.D412–6.

Kanshin E. et al., 2015. A cell-signaling network temporally resolves specific versus promiscuous phosphorylation. Cell reports, 10(7), pp.1202–1214.

Kemmeren P. et al., 2014. Large-scale genetic perturbations reveal regulatory networks and an abundance of gene-specific repressors. Cell, 157(3), pp.740–752.

Machado D. & Herrgard M., 2014. Systematic evaluation of methods for integration of transcriptomic data into constraint-based models of metabolism. PLoS computational biology, 10(4), p.e1003580.

Martin D.E., Soulard A. & Hall M.N., 2004. TOR regulates ribosomal protein gene expression via PKA and the Forkhead transcription factor FHL1. Cell, 119(7), pp.969–979.

Mathelier A. et al., 2014. JASPAR 2014: an extensively expanded and updated open-access database of transcription factor binding profiles. Nucleic acids research, 42(Database issue), pp.D142–7.

Merlini L., Dudin O. & Martin S.G., 2013. Mate and fuse: how yeast cells do it. Open biology, 3(3), p.130008.

Mischnik M. et al., 2016. IKAP: A heuristic framework for inference of kinase activities from Phosphoproteomics data. Bioinformatics, 32(3), pp.424–431.

Mitchell A., Wei P. & Lim W.A., 2015. Oscillatory stress stimulation uncovers an Achilles’ heel of the yeast MAPK signaling network. Science. Available at: http://dx.doi.org/10.1126/science.aab0892.

Mo M.L., Palsson B.O. & Herrgard M.J., 2009. Connecting extracellular metabolomic measurements to intracellular flux states in yeast. BMC systems biology, 3, p.37.

Nielsen P.S. et al., 2001. Transcriptional regulation of the Saccharomyces cerevisiae amino acid permease gene BAP2. Molecular & general genetics: MGG, 264(5), pp.613–622.

O’Duibhir E. et al., 2014. Cell cycle population effects in perturbation studies. Molecular systems biology, 10, p.732.

Oliveira A.P., Ludwig C., et al., 2015. Dynamic phosphoproteomics reveals TORC1-dependent regulation of yeast nucleotide and amino acid biosynthesis. Science signaling, 8(374), p.rs4.

Oliveira A.P., Dimopoulos S., et al., 2015. Inferring causal metabolic signals that regulate the dynamic TORC1-dependent transcriptome. Molecular systems biology, 11(4), p.802.

Oliveira A.P. et al., 2012. Regulation of yeast central metabolism by enzyme phosphorylation. Molecular systems biology, 8. Available at: http://www.nature.com/doifinder/10.1038/msb.2012.55.

Oughtred R., Chatr-Aryamontri A., et al., 2016. BioGRID: A Resource for Studying Biological Interactions in Yeast. Cold Spring Harbor protocols, 2016(1), p.db.top080754.

Oughtred R., Chatr-aryamontri A., et al., 2016. Use of the BioGRID Database for Analysis of Yeast Protein and Genetic Interactions. Cold Spring Harbor protocols, 2016(1), p.db.prot088880.

Park J.O. et al., 2016. Metabolite concentrations, fluxes and free energies imply efficient enzyme usage. Nature chemical biology. Available at: http://dx.doi.org/10.1038/nchembio.2077.

Patil K.R. & Nielsen J., 2005. Uncovering transcriptional regulation of metabolism by using metabolic network topology. Proceedings of the National Academy of Sciences of the United States of America, 102(8), pp.2685–2689.

Pedregosa F. et al., 2011. Scikit-learn: Machine Learning in Python. Journal of machine learning research: JMLR, 12, pp.2825–2830.

Pedruzzi I. et al., 2003. TOR and PKA signaling pathways converge on the protein kinase Rim15 to control entry into G0. Molecular cell, 12(6), pp.1607–1613.

Rhee H.S. & Pugh B.F., 2011. Comprehensive genome-wide protein-DNA interactions detected at single-nucleotide resolution. Cell, 147(6), pp.1408–1419.

Sadowski I. et al., 2013. The PhosphoGRID Saccharomyces cerevisiae protein phosphorylation site database: version 2.0 update. Database: the journal of biological databases and curation, 2013, p.bat026.

Saito H. & Posas F., 2012. Response to hyperosmotic stress. Genetics, 192(2), pp.289–318.

Schacht T. et al., 2014. Estimating the activity of transcription factors by the effect on their target genes. Bioinformatics, 30(17), pp.i401–7.

Schmidt A. et al., 1998. The TOR nutrient signalling pathway phosphorylates NPR1 and inhibits turnover of the tryptophan permease. The EMBO journal, 17(23), pp.6924–6931.

Schulz J.C. et al., 2014. Large-scale functional analysis of the roles of phosphorylation in yeast metabolic pathways. Science signaling, 7(353), p.rs6.

Siddiqui A.H. & Brandriss M.C., 1989. The Saccharomyces cerevisiae PUT3 activator protein associates with proline-specific upstream activation sequences. Molecular and cellular biology, 9(11), pp.4706–4712.

Subramanian A. et al., 2005. Gene set enrichment analysis: a knowledge-based approach for interpreting genome-wide expression profiles. Proceedings of the National Academy of Sciences of the United States of America, 102(43), pp.15545–15550.

Swinnen E. et al., 2006. Rim15 and the crossroads of nutrient signalling pathways in Saccharomyces cerevisiae. Cell division, 1, p.3.

Tachibana C. et al., 2005. Combined global localization analysis and transcriptome data identify genes that are directly coregulated by Adr1 and Cat8. Molecular and cellular biology, 25(6), pp.2138–2146.

Vaga S. et al., 2014. Phosphoproteomic analyses reveal novel cross-modulation mechanisms between two signaling pathways in yeast. Molecular systems biology, 10, p.767.

Venters B.J. et al., 2011. A comprehensive genomic binding map of gene and chromatin regulatory proteins in Saccharomyces. Molecular cell, 41(4), pp.480–492.

Waskom M. et al., 2014. seaborn: v0.5.0 (November 2014), ZENODO. Available at: http://zenodo.org/record/12710.

Wasserman W.W. & Sandelin A., 2004. Applied bioinformatics for the identification of regulatory elements. Nature reviews. Genetics, 5(4), pp.276–287.

Yugi K. et al., 2014. Reconstruction of insulin signal flow from phosphoproteome and metabolome data. Cell reports, 8(4), pp.1171–1183.

Zelezniak A., Sheridan S. & Patil K.R., 2014. Contribution of network connectivity in determining the relationship between gene expression and metabolite concentration changes. PLoS computational biology, 10(4), p.e1003572.

